# Suppression of B-Cell Activation by Human Cord Blood-Derived Stem Cells (CB-SC) through the Galectin-9-Dependent Cell Contact Mechanism

**DOI:** 10.1101/2021.10.07.463564

**Authors:** Wei Hu, Xiang Song, Haibo Yu, Sophia Fan, Andrew Shi, Jingyu Sun, Hongjun Wang, Laura Zhao, Yong Zhao

**Affiliations:** Center for Discovery and Innovation, Hackensack Meridian Health, Nutley, NJ, USA; Throne Biotechnologies, Paramus, NJ, USA; Department of Chemistry and Chemistry Biology, Stevens Institute of Technology, Hoboken, NJ, USA

**Author notes:** **Corresponding author: Yong Zhao M.D., Ph.D.**, Senior Scientist, Throne Biotechnologies, 10 Forest Avenue Suite 110, Paramus, NJ 07652 USA.

**Keywords:** cord blood-derived stem cells, Stem Cell Educator therapy, B cells, Galectin-9, immune modulation, Type 1 Diabetes, autoimmune diseases

## Abstract

**Background:** We developed the Stem Cell Educator therapy among multiple clinical trials based on the immune modulations of multipotent cord blood-derived stem cells (CB-SC) on different compartments of immune cells such as T cells and monocytes/macrophages in diabetes and other autoimmune diseases. However, the effects of CB-SC on the B cells remained unclear. To better understand the molecular mechanisms underlying the immune education of CB-SC, we explored the modulations of CB-SC on human B cells.

**Methods:** CB-SC were isolated from human cord blood units and confirmed by flow cytometry with different markers for their purity. B cells were purified by using anti-CD19 immunomagnetic beads from human peripheral blood mononuclear cells (PBMC). Next, the activated B cells were treated in the presence or absence of coculture with CB-SC for 7 days before undergoing flow cytometry analysis of phenotypic change with different markers. RT-PCR was utilized to evaluate the levels of galectin expressions with or without treatment of activated B cells in order to find the key galectin contributing to the B-cell modulation.

**Results:** Flow cytometry demonstrated that the proliferation of activated B cells was markedly suppressed in the presence of CB-SC, leading to the down-regulation of immunoglobulin productions from the activated B cells. Phenotypic analysis revealed that treatment with CB-SC increased the percentage of IgD^+^CD27^-^ naïve B cells, but decreased the percentage of IgD^-^CD27^+^ switched B cells. Transwell assay showed that the immune suppression of CB-SC on B cells was dependent on the manner of cell-cell contact via Gal-9 molecule, as confirmed by the blocking experiment with the anti-Gal-9 monoclonal antibody. Mechanistic studies demonstrated that both calcium levels of cytoplasm and mitochondria were down-regulated after the treatment with CB-SC, causing the decline of mitochondrial membrane potential in the activated B cells. Western blot exhibited that the levels of phosphorylated Akt and Erk1/2 signaling proteins in the activated B cells were also markedly reduced in the presence of CB-SC.

**Conclusions:** CB-SC displayed multiple immune modulations on B cells through the Gal-9-mediated cell-cell contact mechanism and calcium flux/Akt/Erk1/2 signaling pathways. The data advances current understanding about the molecular mechanisms underlying the Stem Cell Educator therapy to treat autoimmune diseases in clinics.

## Introduction

Human cord blood-derived stem cells (CB-SC) display a unique phenotype with both embryonic and hematopoietic markers that distinguish them from other known types of stem cells, including hematopoietic stem cells (HSC) and mesenchymal stem cells (MSC)[1, 2]. Our previous studies demonstrated that human CB-SC display multiple immune modulations on T cells and monocytes/macrophages via surface molecules and released exosomes[3, 4]. Based on CB-SC’s immunomodulation, we developed the Stem Cell Educator^®^ (SCE) therapy to treat immune dysfunction-associated diseases, including type 1 diabetes (T1D), type 2 diabetes (T2D) and alopecia areata (AA)[5-7], through the multicenter international clinical trials in the United States, China and Spain. SCE therapy circulates a patient’s peripheral blood mononuclear cells (PBMC) through a blood cell separator, cocultures their immune cells with adherent CB-SC *in vitro*, and then returns the “educated” immune cells back to the patient’s blood circulation. Our clinical data demonstrates the safety and clinical efficacy of SCE therapy in reversing the autoimmunity, promoting the regeneration of islet β cells, and improving the metabolic control in T1D and T2D patients[5, 6].

B cells have an important role in maintaining homeostasis and the adaptive immune response through antibody production, antigen presentation and the production of multiple cytokines [8, 9]. Dysfunctions of B cells actively contribute to the pathogenesis of diabetes [10-13] and multiple autoimmune diseases [14, 15]. For example, their roles as antibody-producing cells in systemic lupus erythematosus (SLE)[16] and antigen-presenting cells in T1D and rheumatoid arthritis (RA) have been well recognized [10, 17]. Therefore, it is necessary to correct B cell-associated immune dysfunctions for the treatment of autoimmune diseases. Additionally, galectins are a family of highly-conserved glycan-binding proteins expressed in different tissues, including immune and non-immune cells. In the immune system, galectins are important regulators among innate and adaptive immune responses by regulating a variety of immune cell activations, maturations and other activities. Galectins (Gal)-1,-3, and -9 have shown different effects on the functioning of T cells by modulating their development, activation and differentiation [18-20]. However, the actions of galectins in B cells have only recently begun to be deciphered. Gal-9(Gal-9) is a 34-39 kDa tandem-repeat type protein, which is found in immune cells, endothelial cells, and stem cells [21, 22]. Increasing evidence demonstrated that Gal-9 could not only suppress T-cell activation via the Tim-3 or PD-1 receptor on T cells [23], but also could suppress B-cell activation through the B-cell receptor[24, 25]. To date, our mechanistic studies have demonstrated the immune modulations of SCE therapy on the activated T cells, autoimmune memory T cells [26], regulatory T cells (Tregs) [5] and monocytes/macrophages [4, 6, 27]. The effects of CB-SC on B cells remained elusive. Here, we demonstrated the direct immune modulation of CB-SC on the activated B cells via Gal-9-mediated cell-cell contact mechanism, leading to the marked suppression of B-cell proliferation and phenotypic changes.

## Materials and Methods

### B-cell isolation and culture

Human peripheral blood mononuclear cells (PBMC) (N = 12, aged from 31 to 64 years with average at 46.83 ± 9.67 years old, male =7, female = 5) were isolated by ficoll-hypaque density gradient (GE Healthcare, IL, USA) from the buffy coats purchased from the New York Blood Center (New York, USA). PBMC cell suspensions were pre-treated with anti-CD19-conjugated microbeads (Miltenyi Biotec, CA, USA) according to the instructions of the manufacturer. The purity of positively selected CD19^+^ cells was more than 95%, as assessed by flow cytometry with Korman orange-conjugated mouse anti-human CD19 monoclonal antibody (mAb) (Beckman Coulter, CA, USA). The purified CD19^+^ B cells were cultured in the chemical-defined and serum-free X’VIVO 15 medium (Lonza, Walkersville, MD, USA), in the absence or presence of 100 U/ml penicillin, and 100 μg/ml streptomycin.

### Proliferation Assay

To examine the effects of CB-SC on B-cell proliferation, B cells were stimulated by the following combination at 37°C and 5% CO_2_ conditions: the goat anti-human IgM F (ab’)2 (10 µg/mL), recombinant CD40L (rCD40L 1μg/mL), IL-2 (10 ng/mL), IL-10 (20 ng/mL), and IL-21 (50 ng/mL) in the presence of the treatment with CB-SC for 7 days at the CB-SC:B cells ratios of 1:2, 1:5, and 1:10 in duplicates. The stimulated and unstimulated B cells in the absence of CB-SC served as positive and negative controls respectively. To detect the B-cell proliferation, the purified B cells were initially labeled with carboxyfluorescein succinimidyl ester (CFSE) (Life Technologies, CA, USA), according to the manufacturer protocol. Consequently, the proliferation of B cells was detected by flow cytometry.

### Cell culture for CB-SC

The culture of CB-SC was performed as previously described [4, 5, 27]. In brief, human umbilical cord blood units were collected from healthy donors and purchased from Cryo-Cell international blood bank (Oldsmar, FL, USA). Cryo-cell has received all accreditations for cord blood collections and distributions, with hospital institutional review board (IRB) approval and signed consent forms from donors. Mononuclear cells were isolated with Ficoll-hypaque (γ = 1.077, GE Health) and red blood cells were lysed using ammonium-chloride-potassium (ACK) lysis buffer (Lonza, MD, USA). The remaining mononuclear cells were seeded in 150 x15 mm style non-tissue culture-treated petri dishes or non-tissue culture-treated 24-well plates at 1 ×10^6^ cells/mL. Cells were cultured in X’VIVO 15 chemically-defined serum-free culture medium and incubated at 37°C with 8% CO_2_ for 10-14 days.

### Quantitative Real Time PCR assay

The mRNA expressions of the galectin family were analyzed by quantitative real-time PCR (RT-PCR). Total RNA was extracted from CB-SC using RNeasy mini Kit (Qiagen, CA, USA). First-strand cDNA were synthesized from total RNA using an iScript gDNA Clear cDNA synthesis Kit according to the manufacturer’s instruction (Bio-Rad, Hercules, CA, USA). Real-time PCR was performed using the StepOnePlus Real-time PCR system (Applied Biosystems, CA, USA) under the following conditions: 95 °C for 10 min, then 40 cycles of 95 °C for 15 s, and 60 °C for 60 s. cDNA was amplified with validated specific primers (**Table 1**)[28]. β-actin was used as control.

**Table 1.**
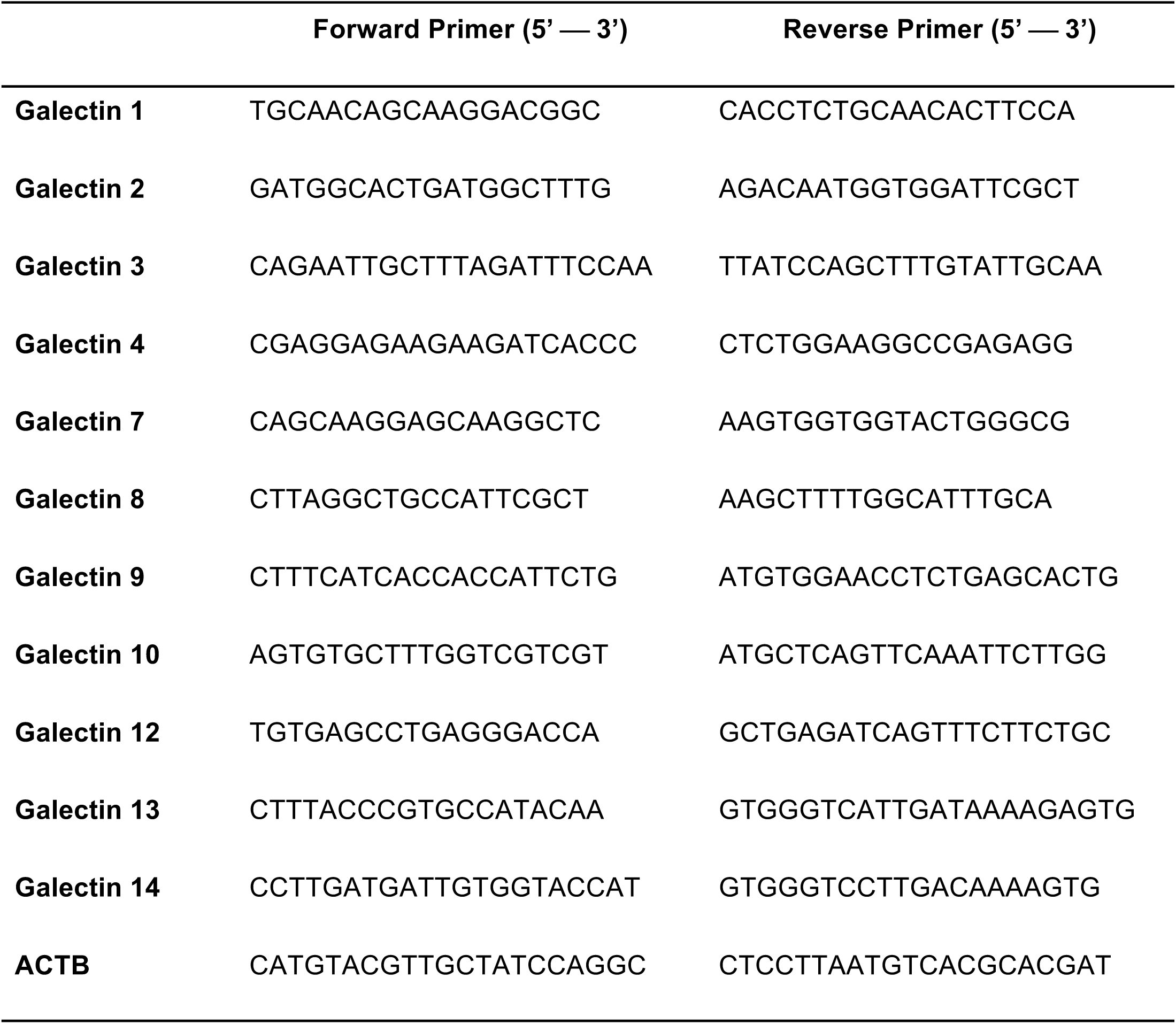
List of primer sequences for RT-PCR analysis of human galectins.

### Assay for antibody production

To detect the antibodies produced by B cells, B cells were stimulated by the following combination: goat anti-human IgM F (ab’)2 (10 µg/mL), recombinant CD40L (rCD40L 1 μg/mL), IL-2 (10 ng/mL), IL-10 (20 ng/mL), and IL-21 (50 ng/mL) in the presence or absence of the treatment with CB-SC in 24-well plate, with 500 µL X’VIVO 15 chemical-defined serum-free culture medium (Lonza, MD, USA) per well. After the treatment for 7 days, the supernatants were collected to determine the levels of antibody productions (e.g., IgG1, IgG2, IgG3, IgG4, IgA, and IgM) by using LEGENDplex^™^ Human Immunoglobulin Isotyping Panel (Biolegend, CA, USA). The Gallios Flow Cytometer was utilized to analyze the data according to the manufacturer’s recommended protocol.

### Blocking Experiments with Gal-9 Antibody

To determine whether Gal-9 contributes to the immune suppression of CB-SC on the activated B cells, the purified CD19 positive B cells were activated with goat anti-human IgM F(ab’)2 (10 µg/mL), recombinant CD40L(rCD40L 1 μg/mL), IL-2(10 ng/mL), IL-10(20 ng/mL), and IL-21(50 ng/mL) in the presence or absence of CB-SC at the ratio of 1:2 in 24-well plate or 6-well plate, with or without adding Gal-9 mAb (10 µg/mL, Biolegend, CA, USA).). The blocking effects of Gal-9 mAb on the B-cell proliferation were examined by flow cytometry with CFSE staining.

To further explore the blocking effects of Gal-9 mAb, the activated B cells were characterized with cytoplasmic and mitochondrial Ca^2+^ levels as well as mitochondrial membrane potential (Δψm) by using flow cytometry as previously described[4]. Briefly, after the 4-hour treatment, B cells were stained with fluorescence dyes including Fluo-4 (ThermoFisher Scientific, MA, USA) for cytoplasmic Ca^2+^, Rhod-2 (ThermoFisher Scientific, MA, USA) for mitochondrial Ca^2+^, and tetramethylrhodamine ethyl ester (TMRE) (Abcam, MA, USA) for detection of mitochondrial membrane potential, respectively.

### Western blot

Cells were prepared with radioimmunoprecipitation assay (RIPA) buffer. Protein concentration was determined by a bicinchoninic acid (BCA) protein assay. The proteins were separated by 10% Tris-HCl gel(Bio-Rad, CA, USA) and transferred to the polyvinylidene fluoride (PVDF) membrane. The proteins were then blotted overnight with anti-human phospho-Akt and anti-human phospho-Erk1/2 mAbs (Cell Signaling, MA, USA), followed by anti-rabbit or anti-mouse horseradish peroxidase(HRP)-conjugated secondary mAb (ThermoFisher scientific) [4]. The membrane was incubated with the chemiluminescent substrate (ThermoFisher Scientifc, CA, USA) and chemiluminescent signal was detected by using ChemiDoc Imaging System (Bio-Rad, CA, USA). β-actin served as an internal control.

### Flow cytometry

Phenotypic characterization of B-cell subsets was performed by flow cytometry [4, 27] with specific markers including PE-conjugated mouse anti-human CD27 (Biolegend, CA, USA) and APC-conjugated mouse anti-human IgD (Biolegend, CA, USA). To determine the purity of CB-SC, CB-SC were examined by flow cytometry with CB-SC-associated markers, including PE-Cy7-conjugated mouse anti-human CD45 (Beckman Coulter, CA, USA), efluor660-conjugated rat anti-human OCT3/4 (ThermoFisher Scientific, MA, USA), FITC-conjugated mouse anti-human SOX2 (ThermoFisher Scientific, MA, USA), BV421-conjugated mouse anti-human CD34 (Biolegend, CA, USA), PE-conjugated mouse anti-human CD270 (ThermoFisher Scientific, MA, USA), and Pacific Blue-conjugated mouse anti-human CD274 (ThermoFisher Scientific, MA, USA) mAbs. Isotype-matched immunoglobulin (IgGs) served as controls.

### Statistical Analysis

Statistical analysis of data was performed with GraphPad Prism 8 (version 8.0.1) software. The normality test of samples was evaluated using the Shapiro-Wilk test. Statistical analysis of data was performed using the two-tailed paired student’s t-test to determine statistical significance for parametric data between untreated and treated groups. The Mann-Whitney U test was utilized for non-parametric data. Values were given as mean ± SD (standard deviation). Statistical significance was defined as *P* < 0.05.

## Results

### CB-SC suppressed the proliferation of activated B cells

Initially, the purity of CB-SC was characterized by flow cytometry with CB-SC-associated markers including leukocyte common antigen CD45, embryonic stem (ES) cell markers OCT3/4 and SOX2, hematopoietic stem cell marker CD34, and immune tolerance-related markers CD270 and CD274. Findings revealed CB-SC highly expressed CD45, OCT3/4, SOX2, CD270 and CD274, but did not express CD34 **(Figure 1A)**. CD45 and Oct3/4 are routinely utilized for the purity test of CB-SC, at ≥ 95% of CD45^+^ OCT3/4^+^ CB-SC.

**Fig.1.**
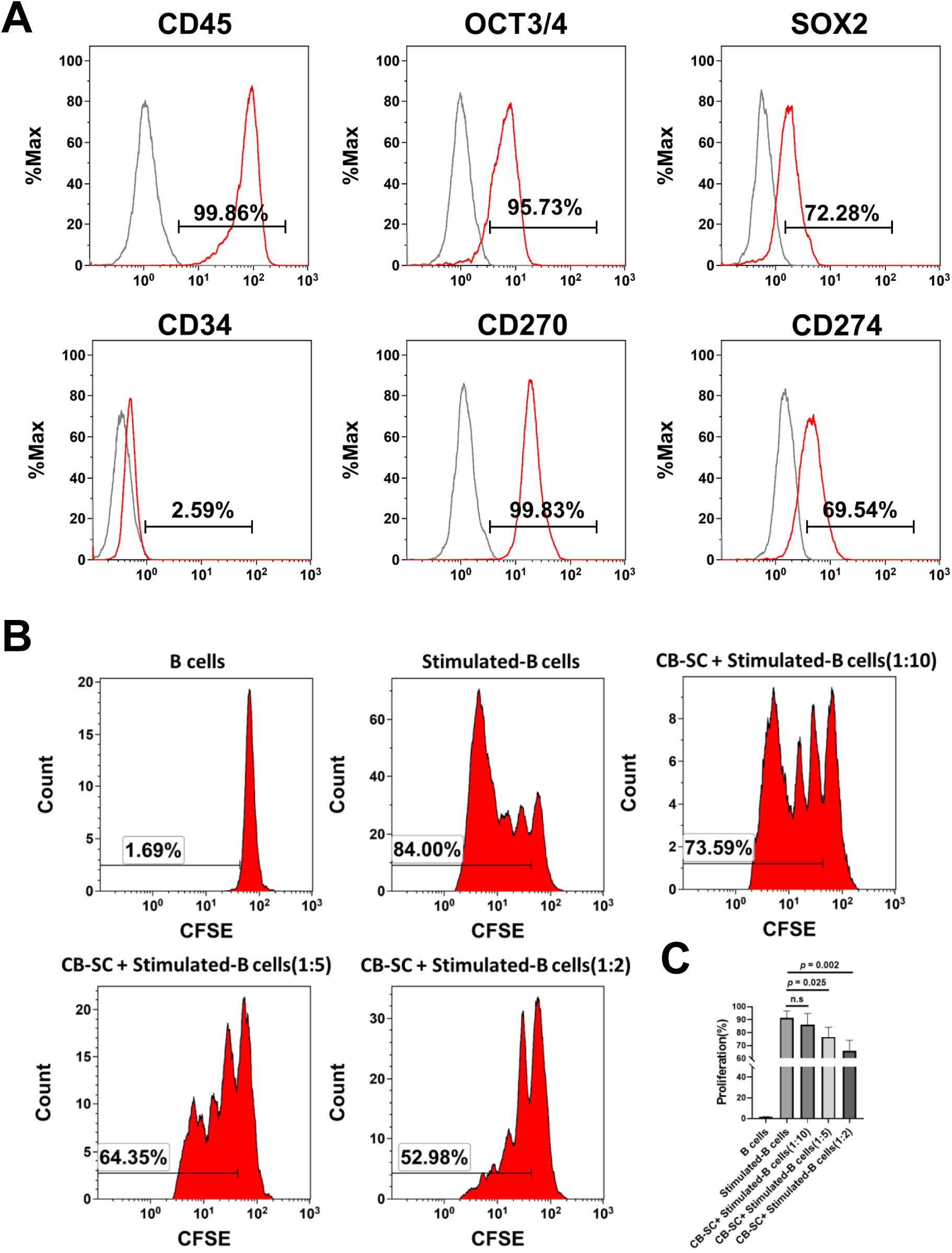
Immunosuppression of Cord blood-derived stem cell on the B cell. **(A)** Phenotypic characterization of CB-SC with high purity. CB-SC were analyzed by flow cytometry with associated markers, including leukocyte common antigen CD45, embryonic stem (ES) cell markers OCT3/4 and SOX2, hematopoietic stem cell marker CD34, and the immune modulation-related marker CD270 and CD274. Isotype matched immunoglobulin G (IgGs) served as control. Data were represented from four experiments with similar results. **(B)** Suppression of B-cell proliferation by CB-SC. The carboxyfluorescein succinimidyl ester (CFSE)-labeled B cells were stimulated to proliferate with activation cocktails in the presence of different ratios of CB-SC. Untreated B cells served as negative control. Histograms of flow cytometry were representative of five experiments with similar results. **(C)** Quantitative analysis of B-cell proliferation shows a remarkable decrease in B cell expansion after the treatment with CB-SC at the different ratios of CB-SC:B cells of 1:10 (n.s, *P* > 0.05, N=6), 1:5 (*P* = 0.025, N=6), and 1:2 (*P* = 0.002, N=6)

To explore the immune modulation of CB-SC on B cells, CB-SC were co-cultured with B cells at different ratios of CB-SC. B cells (e.g., 1:2, 1:5, and 1:10) were activated in the presence of cocktails (anti-IgM, rCD40L, IL-2, IL-4, IL-10, IL-21)[24]. The proliferation of B cells was examined by flow cytometry, after carboxyfluorescein succinimidyl (CFSE) staining and in combination with propidium iodide (PI) staining, to determine the dead cells. The results demonstrated that there were no differences in the percentages of dead cells among different groups of CB-SC treatments relative to that of CB-SC-untreated B cells **(Figure S1)**. However, the percentages of B-cell proliferation markedly declined from 91.52± 5.31 to 65.84 ± 8.24 at the ratio of 1:2 (*p* < 0.005), and 76.78 ± 7.44 at the ratio of 1:5 (*p* < 0.05), respectively **(Figure 1B and C)**. The data indicated the suppression of CB-SC on the B-cell proliferation.

### CB-SC inhibited immunoglobulin production

Immunoglobulin (Ig) are proteins secreted by plasma B cells and are present on the surface of B cells (e.g., IgD). They are assembled from identical couples of heavy (H) and light (L) chains. Based on the difference among heavy chains, immunoglobulins are characterized by 5 classes of Ig including IgM, IgG, IgA, IgE, and IgD. To explore the effects of CB-SC on B cells, Ig productions were examined by using flow cytometry. Comparing with Ig productions of CB-SC-untreated B cells, the data showed that CB-SC could markedly inhibit Ig productions at the ratio of CB-SC:B-cells at 1:5, including IgG1 production (*p* = 0.019) **(Figure 2A)**, IgG2 (*p* = 0.036) **(Figure 2B)**, IgG3 (*p* = 0.037) **(Figure 2C)**, IgG4 (*p* = 0.037) **(Figure 2D)**, IgA (*p* = 0.008) **(Figure 2E)**, and IgM (p=0.024) **(Figure 2F)**.

**Fig.2.**
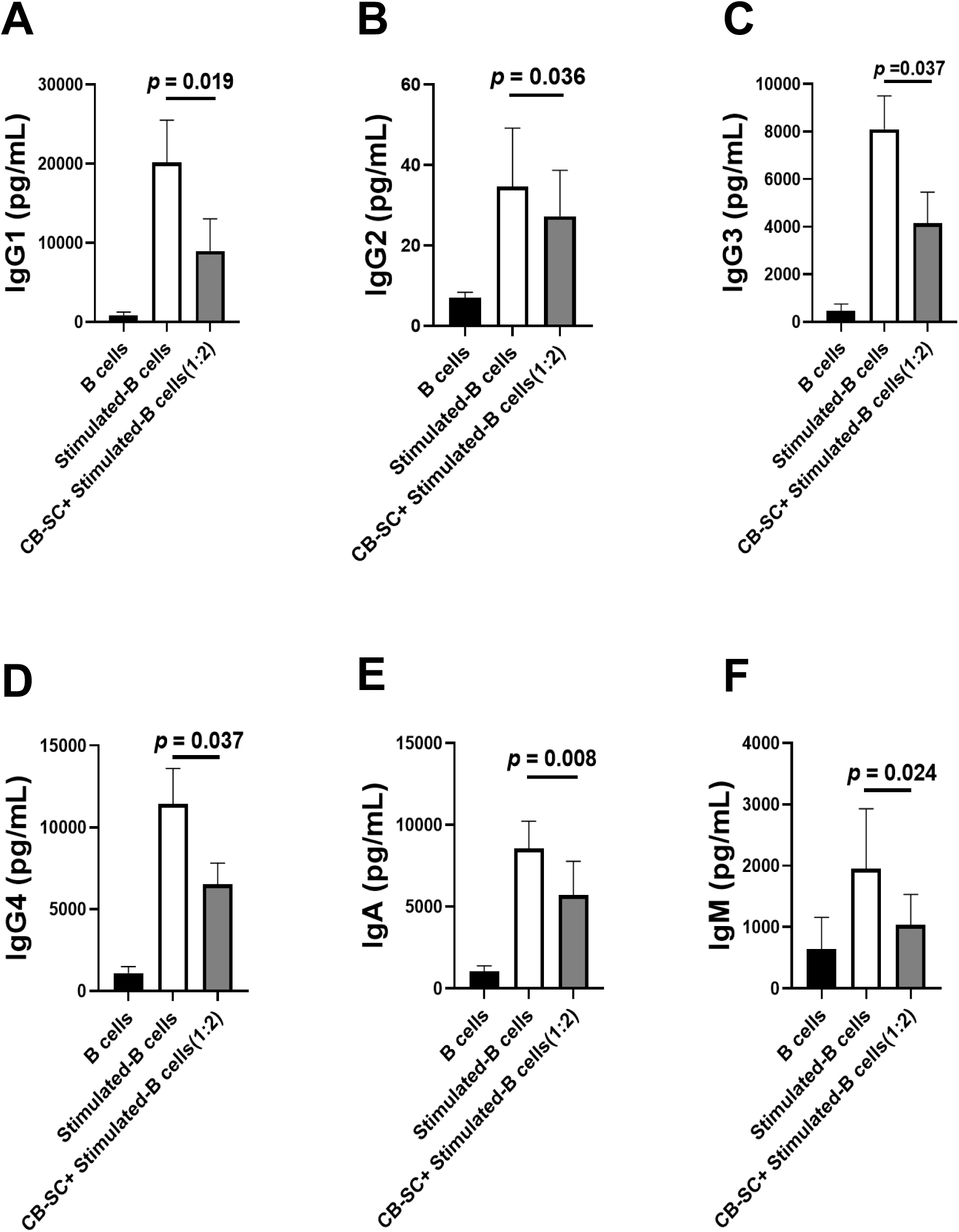
Inhibition of B-cell immunoglobulin production by CB-SC. B cells were stimulated in the presence of cocktails (anti-IgM, rCD40L, IL-2, IL-4, IL-10, IL-21). CB-SC markedly inhibit Ig productions of stimulated B cells at the ratio of CB-SC : B-cells at 1:5. Untreated B cells served as negative control. **(A)** CB-SC inhibit IgG1 production. *P* = 0.019 (N=4). **(B)** CB-SC inhibit IgG2 production. *P* = 0.036 (N=6). **(C)** CB-SC inhibit IgG3 production. *P* = 0.037 (N=4). **(D)** CB-SC inhibit IgG4 production. p=0.037 (N=6). **(E)** CB-SC inhibit IgA production. *P* = 0.008 (N=4). **(F)** CB-SC inhibit IgM production. *P* = 0.024 (N=4).

### Modulation of CB-SC on naïve and memory B cells

CD27 and IgD are widely accepted biomarkers used to characterize B cells into memory and naive subsets, such as naïve B cells (CD27^-^IgD^+^), switched memory B cells (CD27^+^IgD^-^), and non-switched memory B cells (CD27^+^ IgD^+^) [29]. To explore the action of CB-SC on memory B cells, the activated B cells were treated with or without CB-SC. Flow cytometry established that both percentages of naïve B cells and switched B cells were significantly downregulated after the B-cell activation. However, their percentages were markedly changed after the treatment with CB-SC (**Figure 3A, B and D**). The percentage of naïve B cells was increased from 8.79 ± 2.14 for CB-SC-untreated B cells to 28.23 ± 4.82 for CB-SC-treated B cells at the ratio of 1:2. (*P* = 0.003, **Figure 3B**). The percentage of switched memory B cells was decreased from 63.53 ± 6.85 for CB-SC-untreated B cells to 48.75 ± 3.09 for CB-SC-treated B cells at the ratio of 1:2. (*P* = 0.0023, **Figure 3D**). Notably, the percentage of non-switched CD27^+^IgD^+^ memory B cells failed to mark changes before and after the treatment with CB-SC (**Figure 3C**). The data suggest the modulation of CB-SC on the B-cell differentiation.

**Fig.3.**
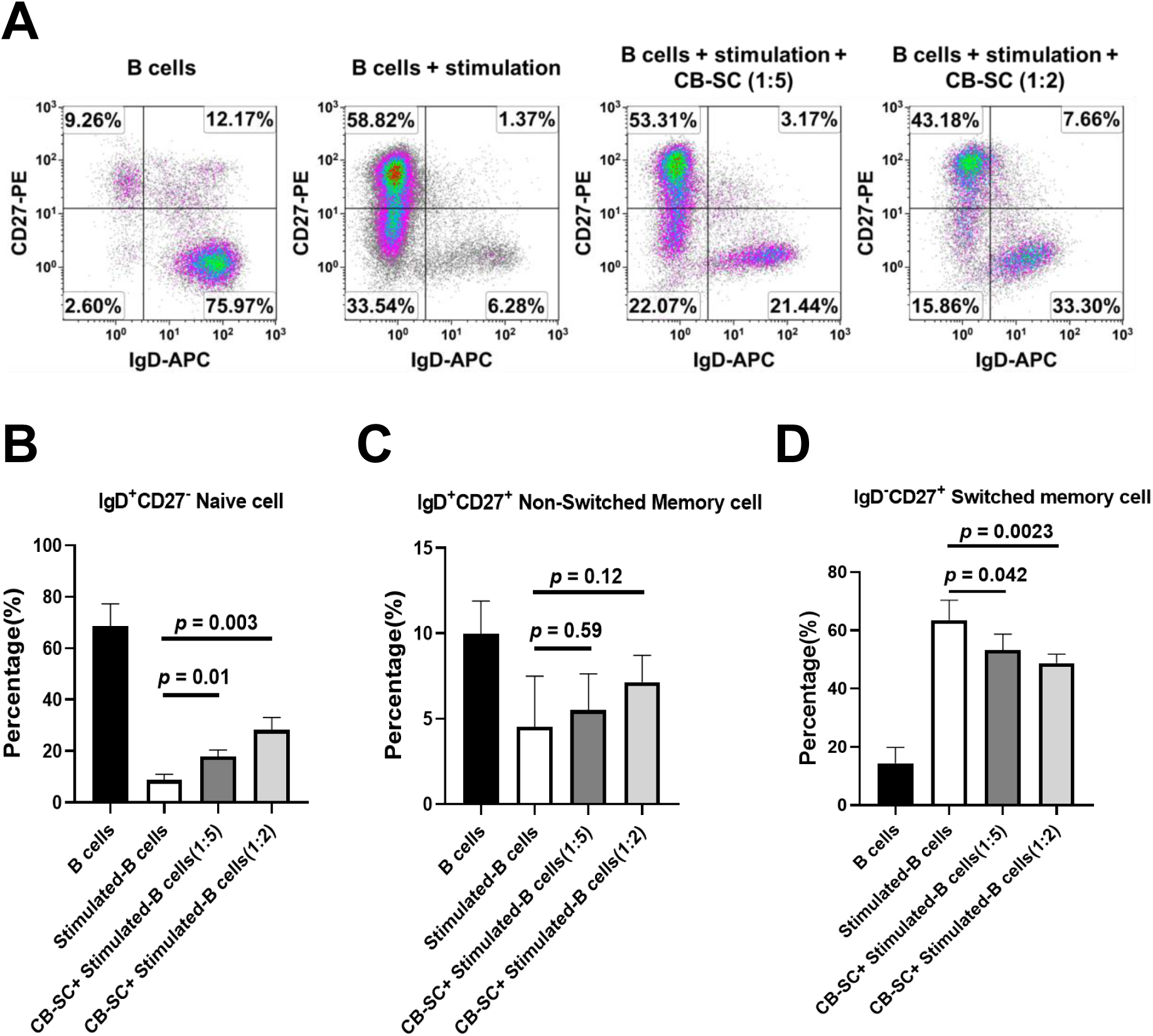
Modulation of different B cell subpopulation by CB-SC. **(A)** Upregulation of the percentage of naïve B cells by CB-SC and downregulation of the percentage of switched B cells. Histograms of flow cytometry were representative of four experiments with similar results. Isotype-matched IgGs served as negative controls. **(B)** Increase in the percentage of naïve B cells after the treatment with CB-SC at ratio of 1:5 (*P* = 0.01, N = 4), 1:2 (*P* = 0.003, N = 4). **(C)** There is no significant effect on the percentage of non-switched B cells. **(D)** Decrease in the percentage of switched B cells. Results were given as mean ± SD. *P* < 0.05 as a significant difference.

### CB-SC mediate B cell suppression by cell-cell contact

Our previous studies demonstrated that multiple mechanisms contribute to the immune modulations of CB-SC on T cells, such as PD-L1/PD1-mediated cell-cell inhibition and releasing soluble factors (e.g., nitric oxide and transforming growth factor-β1) [30]. To clarify whether cell-cell contact or soluble factors were involved in the immune modulations of CB-SC on B cells, we performed transwell experiments. Flow cytometry demonstrated that the suppression of CB-SC on B-cell proliferation was abolished with the transwell co-culture system (**Figure 4A and B**), highlighting how the surface molecule expressed on CB-SC contributes to the suppression via cell-cell contact mechanism.

**Fig 4:**
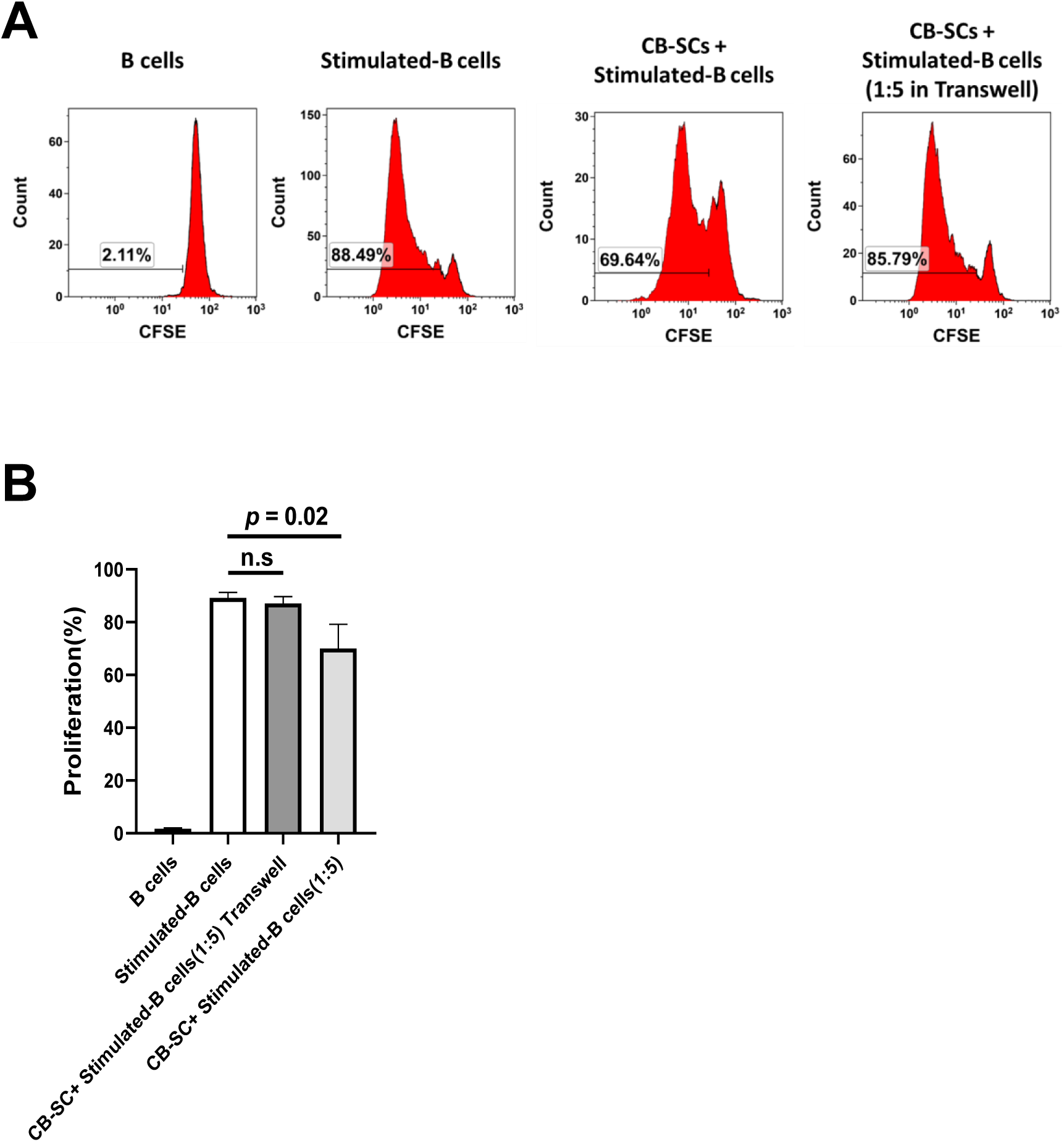
Cell-cell contact mechanism contributes to the CB-SC-mediated immune suppression. **(A)** CB-SC cocultured with CFSE-labeled stimulated B cells in transwells. Both unstimulated and stimulated B cells without CB-SC coculture served as control. Histograms of flow cytometry were representative of four experiments with similar results. **(B)** CB-SC failed to suppress the proliferation of stimulated B cells in transwell coculture system at the ratio of CB-SC:B cells of 1:5. In contrast, the direct coculture of CB-SC with stimulated B cells displayed the marked inhibition of B-cell proliferation at the same ratio (*P* = 0.02, N=4).

### Expression of Gal-9 on CB-SC acts as a key molecule contributing to the B-cell modulation

Galectins have been recognized as essential regulators contributing to the induction of an immune tolerance and homeostasis, therefore functioning as the attractive therapeutic targets for attenuating autoimmune and inflammation disorders [20, 31]. To determine whether galectins were involved in the immune modulation of CB-SC on B cells, we initially analyzed the profile of galectin mRNA expressions including Gal-1, -2, -3, -4, -7, -8, -9, -10, -12, -13, and -14 **(Figure 5A)**. RT-PCR results showed that there were higher expressions of Gal-1, -2, -3, -4, -7, -8, and -9 mRNA than those of others (e.g., Gal-10, - 12, -13, and -14) in CB-SC **(Figure 5A)**. In comparison with the changes of other galectin mRNA levels, the level of Gal-9 mRNA expression was markedly increased approximately 10-fold after being co-cultured with the activated B cells for 48 hours **(Figure 5B)**. To confirm whether the protein level of Gal-9 was upregulated, we performed flow cytometry with Gal-9 mAb. The flow data revealed that the median fluorescence intensity (MFI) in CB-SC was significantly increased in the presence of activated B cells **(Figure 5C, 5D)**. To further prove the involvement of Gal-9 in the immune modulation of CB-SC on B cells, we performed the blocking experiment with neutralizing Gal-9 mAb. The data demonstrated that the suppression of CB-SC on the proliferation of activated B cells was markedly reversed after the treatment with the neutralizing Gal-9 mAb relative to that of the Gal-9 mAb-untreated group (*P* = 0.025, **Figure 5E**). This finding indicated that Gal-9 contributed to the immune modulation of CB-SC on activated B cells.

**Fig. 5.**
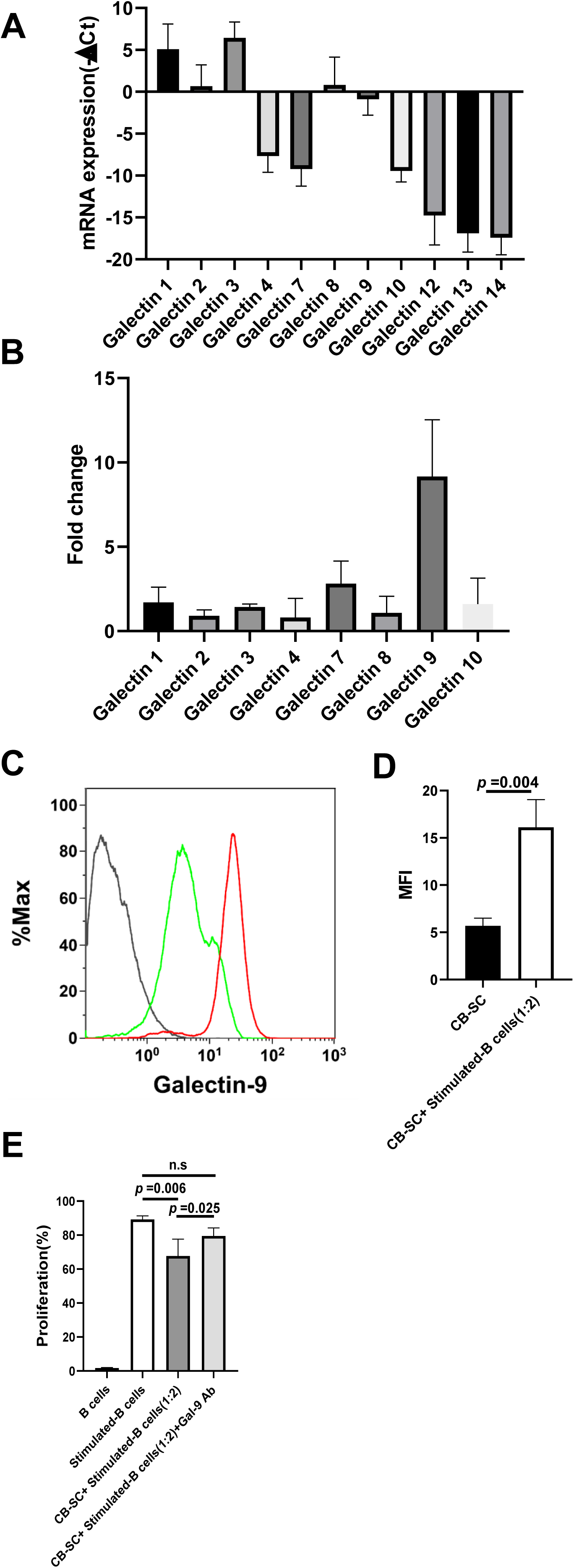
Gal-9 expressed on CB-SC acts as the key mediator for the CB-SC-induced B-cell suppression. **(A)** RT-PCR analysis of galectin expressions in CB-SC. Δ actin served as control. **(B)** Changes in the levels of galectin expressions after coculture of CB-SC with the stimulated B cells for 2 days (N = 3). **(C)** Upregulate the level of Gal-9 expression on CB-SC after coculture with stimulated B cells (red line) relative to that on the untreated CB-SC (green line). Isotype-matched IgG served as negative control. **(D)** Increase the level of Gal-9 expression (median fluorescence intensity, MFI) on CB-SC after coculture with stimulated B cells. The data were given as mean ± SD of three experiments with CB-SC (N = 3)-treated B cells (N = 3). **(E)** Stimulated B cells were coculture with CB-SC at ratio of 1:2 in the presence or absence of Gal-9 mAb blocking. The inhibition of CB-SC on B-cell proliferation was abolished after blocking with Gal-9 mAb (stimulated B cells with CB-SC *vs* stimulated B cells + CB-SC + Gal-9 mAb, *P* = 0.025, N=4).

### Suppression calcium flux by CB-SC was Gal-9 dependent

The activation and proliferation of B cells are initiated by the B cell receptor (BCR), which triggers a number of signaling cascades [32]. The increase in intracellular Ca^2+^ levels is one of the critical signaling pathways for tuning B-cell responses and development post the BCR activation [33]. To further explore the molecular mechanism underlying the inhibition of B-cell proliferation by the treatment with CB-SC, we examined the changes of cytosolic and mitochondrial Ca^2+^ levels in CB-SC-treated B cells by flow cytometry after being stimulated with B cell-dependent activation cocktails. Using the Fluor-4 staining for cytosolic calcium, the median fluorescence intensity of Fluo-4^+^ activated B cells was markedly downregulated in the presence of CB-SC at the ratio of 1:5 (*P* < 0.05). The suppressive effect on cytosolic Ca^2+^ levels was reversed after the blocking with Gal-9 mAb (**Figure 6A**). The direct effect of Gal-9 on the changes of cytosolic Ca^2+^ levels was further confirmed by the treatment with recombinant Gal-9 at 0.5 μg/mL (**Figure 6A**). Using the Rhod-2 staining as an indicator for the mitochondrial calcium, flow cytometry demonstrated the mitochondrial Ca^2+^ levels in the stimulated B cells were significantly reduced after the treatment with CB-SC at the ratio 1:5 of CB-SC to B cells (**Figure 6B**). Similar to the changes of cytosolic Ca^2+^ levels, the mitochondrial Ca^2+^ levels in the CB-SC-treated B cells were increased in the presence of Gal-9 mAb (**Figure 6B**). Mitochondrial membrane potential (Δψm) was also decreased in the stimulated B cells after the treatment with CB-SC, but upregulated after the blocking with Gal-9 mAb (**Figure 6C**). The data indicated that the Gal-9-mediated Ca^2+^ signaling pathway contributed to the modulation of CB-SC on the activated B cells.

**Fig. 6.**
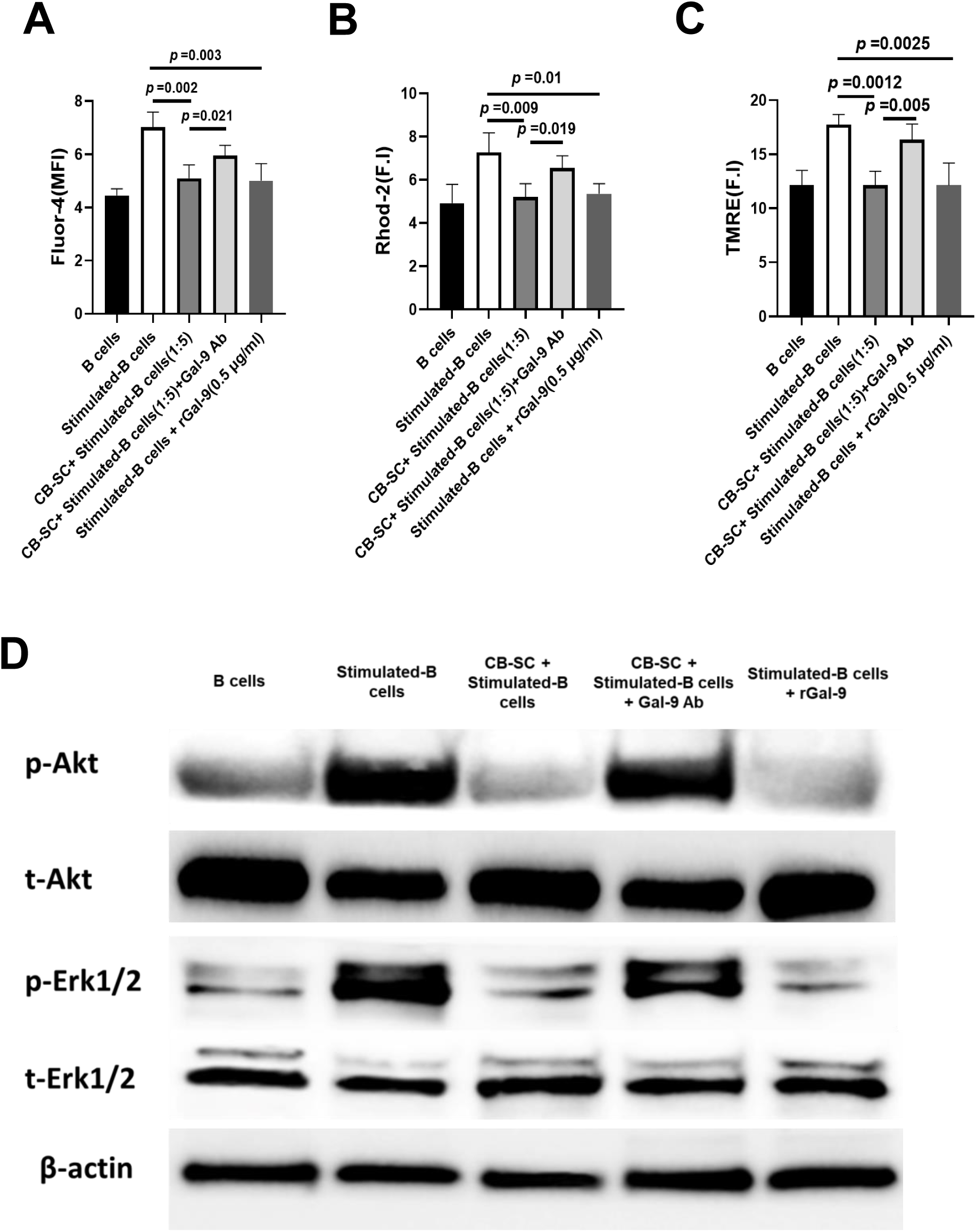
Gal-9 expressed on CB-SC contributes to the modulation of Ca^2+^-associated signaling pathways in stimulated B cells. **(A)** Flow cytometry analysis of cytoplasmic Ca^2+^ with Fluor-4 staining shows the remarkable decrease in the median fluorescence intensity (MFI) value of Fluor-4^+^ B cells after the treatment with CB-SC (*P* < 0.05) or 0.5 μg/mL rGal-9 (*P* < 0.01), but markedly increased after blocking with Gal-9 mAb. **(B)** Flow cytometry analysis of mitochondrial Ca^2+^ with Rhod-2 staining shows the substantial decline in the MFI value of Rhod-2^+^ B cells after the treatment with CB-SC (*P* < 0.01) or 0.5 μg/mL rGal-9 (*P* < 0.01), but clearly improved after blocking with Gal-9 mAb. **(C)** Downregulate the mitochondrial membrane potential in the stimulated B cells after the treatment with CB-SC, which was in the Gal-9-dependent manner as demonstrated by flow cytometry analysis after staining with tetramethylrhodamine, ethyl ester (TMRE). **(D)** Western blotting showed the reduced expression of phosphorylated AKT and ERK1/2 in stimulated B cells after the treatment with CB-SC or rGal-9, but upregulated after blocking with Gal-9 mAb. β-actin served as control.

Additionally, Western blotting showed that both BCR downstream molecules phospho-Akt and phospho-Erk1/2 were upregulated in the stimulated B cells without affecting their total protein levels, but markedly downregulated after the treatment with CB-SC or recombinant human Gal-9 (0.5 µg/mL). Such inhibitory effects of CB-SC on the phospho-Akt and phospho-Erk1/2 were decreased after blocking with Gal-9 mAb (**Figure 6D**). These data suggest that Gal-9 expressed on CB-SC contributed to the immune modulation of CB-SC on B cells via the regulation of Ca^2+^ flux and phosphorylation of Akt and Erk1/2 signaling pathways.

## Discussion

Over the last 10 years, CB-SC have been utilized in multicenter international clinical trials and designated to Stem Cell Educator^®^ (SCE) therapy for the treatment of autoimmune disease including type 1 diabetes (T1D) [5, 26], alopecia areata (AA) [7], and other chronic metabolic inflammation-associated diseases (e.g., type 2 diabetes [6]). Mechanistic studies demonstrated that CB-SC displayed strong immune modulation on T cells and monocytes such as inhibition of T-cell activation and proliferation, percentage reductions of effector memory T cells (T_EM_) [26] and induction of the differentiation of monocytes into anti-inflammation type 2 macrophages (M2) [4, 27]. The modulation of CB-SC on B cells remains unclear. Here, we demonstrated the immunomodulation capabilities of CB-SC on activated B cells by inhibiting the B-cell proliferation and the activation of naïve B cells, down-regulating the differentiation of switched memory B cells, and reducing the production of immunoglobulins. These findings advance our understanding about the molecular mechanism of Stem Cell Educator therapy for the treatment of T1D and other autoimmune diseases.

B cells are important effector cells involved in the pathogenesis of autoimmune diseases through the production of autoantibodies, the promotion of CD4^+^ T cell responses via antigen presentation, and the release of inflammatory cytokines (e.g., TNF-α and IL-6) [15, 34]. Increasing evidence indicates the importance of B cell-mediated autoimmunity in the pathogenesis of T1D, even though T cells are generally considered the major pathogenic effector cells contributing to the destruction of islet β cells. Researchers found that blocking B cells or impairing B cell function will significantly decrease the incidence of diabetes in NOD mice [35, 36]. Additionally, the depletion of B cells with anti-human CD20 antibody (Rituximab) markedly preserved islet β-cell function and improved C-peptide levels after 1 year follow-up in recent-onset T1D patients [37]. The current study demonstrated that CB-SC markedly suppressed the proliferation of activated B cells and reduced the antibody productions in these activated B cells. Therefore, these data suggest the clinical translational potential of Stem Cell Educator therapy to treat other B cell-mediated autoimmune diseases.

To date, the characterization of B-cell phenotype with multiparameter flow cytometry has identified several B-cell subpopulations including CD27^+^IgD^+^ non-switched memory B cells and CD27^+^IgD^-^ switched memory B cells, which may represent a biomarker for some autoimmune diseases. For instance, the percentage of switched memory B cells increased in systemic lupus erythematosus (SLE) and rheumatoid arthritis (RA) [38-40].

Our flow cytometry analysis substantiated that the percentage of switched memory B cells was markedly reduced after the treatment with CB-SC in a dose-dependent manner. Notably, the percentage of naïve B cells (CD27^-^IgD^+^) was increased, highlighting the modulation of CB-SC on B-cell differentiation with the reduction of memory B cells.

To elucidate the molecular basis of SCE therapy, previous studies have identified several molecular and cellular pathways that alter autoimmune T cells and the functions of pathogenic monocytes/macrophages (Mo/Mϕs) to elicit immune tolerance via: (1) the expression of <u>a</u>utoimmune regulator (AIRE) in CB-SC, which is a master transcriptional regulator that acts to eliminate the self-antigen reactive T cells in the thymus and is controlled by the activation of the receptor activator of NF-κB (RANK) signaling pathway [41]; (2) secretion of CB-SC-derived exosomes (cbExosomes), which polarize human blood Mo/Mϕ into type 2 macrophages (M2) [4, 27], further contributing to immune tolerance and preventing β-cell destruction; and (3) migration of platelet-derived mitochondria (pMitochondria) to islets, which are absorbed by pancreatic islets and contribute to an improved proliferation of human islet Δ cells [42].

Our current studies revealed the direct immune modulation of CB-SC on activated B cells through the expression of Gal-9 on CB-SC, as demonstrated by trans-well coculture and blocking experiment with anti-Gal-9 mAb. What’s more, further mechanistic studies confirmed that Gal-9 expressed on CB-SC directly contributed to the regulation of Ca^2+^ flux and phosphorylation of Akt and Erk1/2 signaling pathways in the stimulated B cells. Waters and colleagues reported an increase in the oxidative phosphorylation and mitochondrial membrane potential (Δψm) among the stimulated B cells [43], which was consistent with our current data showing the enhanced median fluorescence intensity of TMRE staining. Notably, the Δψm of stimulated B cells was substantially reduced in the Gal-9-dependent manner.

Galectins are β-galcotosid-bind lectins which can be expressed by different types of stem cells and act as regulators of immune cell function [44], especially galectin-3 and Gal-9. Galectin-3 suppresses the activation of TCR-mediated signal transduction [45], while Gal-9 binds T cell Ig mucin-3 (Tim-3) and induces negative regulate T helper 1(Th1) immunity [46]. Our current data confirmed that galectin-1, 2, 3, 4, 7, 8, and 9 were highly expressed on CB-SC, but only Gal-9 primarily contributed to the immune modulation of CB-SC on activated B cells. Gal-9 was not only located on the cellular membrane, but also acted as a soluble factor involved in the immune modulation [23]. To test this possibility, we found that CB-SC-released Gal-9 was less than 10% of total CB-SC-derived Gal-9 after 3 days culture (**Figure S2**). Therefore, Gal-9 expressed on CB-SC’s membrane displayed more potential than the soluble form of CB-SC-secreted Gal-9 during the B-cell immune modulation. Giovannone et al reported that Gal-9 can directly bind to the poly-LacNAc-containing N-glycans on leukocyte common antigen CD45 of B cells, leading to the diminished intracellular calcium levels and ultimately inhibiting B cell activation [24]. This report was consistent with the reduction of cytosolic Ca^2+^ levels in our current study. Additionally, several studies revealed the distribution of IgM on B-cell surface membranes which form the nanoscale clusters and act as BCR of primary B cells [25, 47, 48]. Using dual-color direct stochastic optical reconstruction microscopy (dSTORM), Cao and colleagues confirmed that Gal-9 can also directly bind to IgM-BCR of murine B cells [25], nearly resulting in a complete abolishment of the BCR activation [25]. Due to the BCR-mediated Ca^2+^ influx as the critical signal for B cell activation [49], the immune modulation of CB-SC on activated B cells primarily targets the regulation of intracellular Ca^2+^ levels through the Gal-9-mediated pathway, leading to dampened B-cell responses and shaping their differentiation. These novel molecular mechanisms will facilitate the clinical translation of Stem Cell Educator therapy to treat T1D and other autoimmune diseases.

## Conclusions

Stem Cell Educator therapy has been unutilized to treat multiple autoimmune- and inflammation-associated diseases, which pathogenesis involve in T cells, B cells, and monocytes/macrophages. The current study revealed that CB-SC displayed multiple immune modulations on B-cell proliferation and differentiation and antibody productions through the Gal-9-mediated cell-cell contact mechanism and calcium flux/Akt/Erk1/2 signaling pathways. These findings lead to a better understanding of the molecular mechanisms of Stem Cell Educator therapy to treat autoimmune diseases in clinics.

## List of abbreviations

AA: alopecia areata
BCR: B cell receptor
CB-SC: cord blood-derived stem cells
CFSE: carboxyfluorescein succinimidyl ester
Gal-9: galectin-9
Ig: Immunoglobulin
MFI: median fluorescence intensity
NOD: non-obese diabetic
PBMC: peripheral blood mononuclear cells
RT-PCR: reverse transcription-polymerase chain reaction
T1D: type 1 diabetes
T2D: type 2 diabetes.

## Acknowledgments

We are grateful to Ludwig for generous funding support via Hackensack UMC Foundation.

## Author Contributions

Y.Z.: supervised experiments, and contributed to concepts, experimental design, data analysis and interpretation, manuscript writing, and final approval of manuscript; W.H.: performed most experiments, contributed to experimental design, and data analysis; H.Y., and X.S., J.S.: isolate B cells and flow cytometry and data analysis. S.F, A.S., L.Z, H.W.: Revised manuscript and English editing. All authors have read and agreed to the published version of the manuscripts.

## Funding

This research received no external funding.

## Availability of data and materials

The data that support the findings of this study are available from the corresponding author upon request.

## Declarations

### Ethics approval and consent to participate

Human cord blood units and buffy coats blood were purchased from Cryo-Cell International blood bank and the New York Blood Center (NYBC), respectively. Both Cryo-Cell and NYBC have received all accreditations for blood collections and distributions, with institutional review board (IRB) approval and signed consent forms from donors.

### Consent for publication

Not applicable.

### Competing Interests

Dr. Yong Zhao was inventor of Stem Cell Educator technology and has a fiducial role at Throne Biotechnologies. All other authors have no financial interests that may be relevant to the submitted work.

## Supplementary data

**Figure S1:**
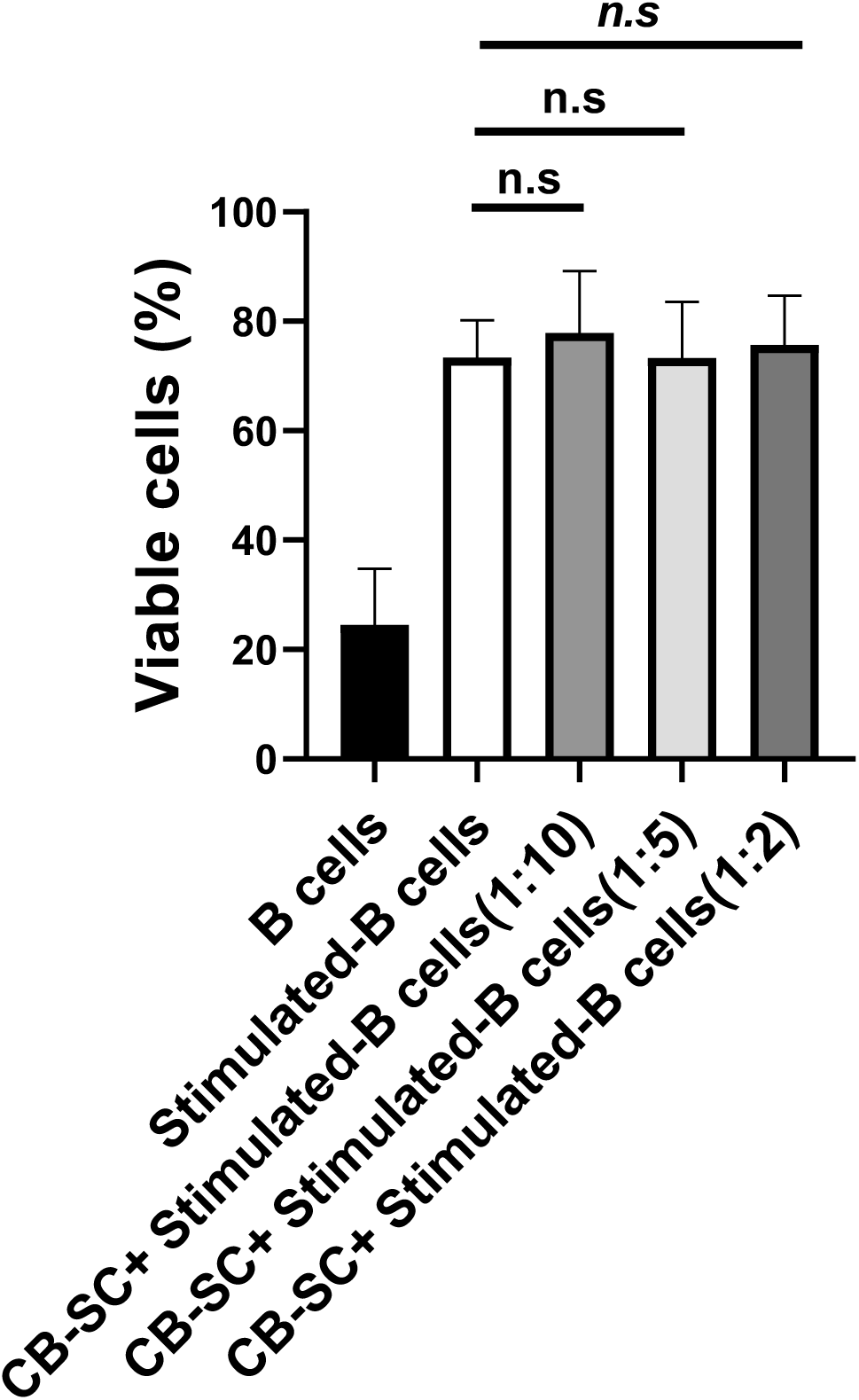
Flow cytometry showed that there were no marked differences on the percentages of viable cells after the treatment with different ratios of CB-SC:B cells, relative to that of stimulated B cells. The viable cells from different samples were gated for analysis after excluding the propidium iodide (PI)-positive dead cells. The data were given as mean ± SD of six experiments with CB-SC (N = 4)-treated B cells (N = 6).

**Figure S2:**
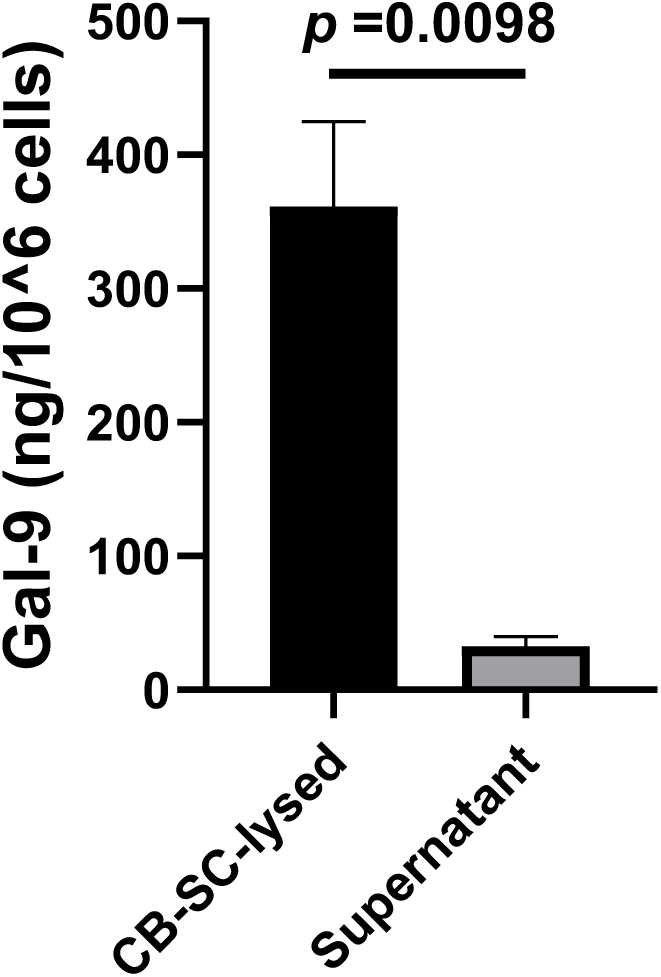
Compare the levels of galectin-9 expressed on CB-SC and released in the supernatants of CB-SC cultures. CB-SC (1 × 10^7^ cells/dish) were cultured in fresh chemical-defined and serum-free X’VIVO 15 medium (Lonza, Walkersville, MD, USA) at 37°C and 8% CO_2_ conditions. After culture for 3 days, CB-SC cells and supernatants were collected respectively for analysis Gal-9 protein concentration. The data were given as mean ± SD of three experiments

